# Novel regulatory variant in ABO intronic RUNX1 binding site inducing A_3_ phenotype

**DOI:** 10.1101/2023.05.04.539366

**Authors:** Gian Andri Thun, Morgan Gueuning, Sonja Sigurdardottir, Eduardo Meyer, Elise Gourri, Linda Schneider, Yvonne Merki, Nadine Trost, Kathrin Neuenschwander, Charlotte Engström, Beat M. Frey, Stefan Meyer, Maja P. Mattle-Greminger

**Author notes:** **Correspondence** Maja P. Mattle-Greminger, Department of Research and Development, Blood Transfusion Service Zurich, Swiss Red Cross, Rütistrasse 19, 8952 Schlieren, Switzerland.

## Abstract

**Background and Objectives:** Mixed-field agglutination in ABO phenotyping (A_3_, B_3_) has been linked to genetically different blood cell populations like in chimerism, or to rare variants in either *ABO* exon 7 or regulatory regions. Clarification of such cases is challenging and would greatly benefit from sequencing technologies that allow resolving full-gene haplotypes at high resolution.

**Materials and Methods:** We used long-read sequencing by Oxford Nanopore Technologies to sequence the entire *ABO* gene, amplified in two overlapping long-range PCR fragments, in a blood donor presented with A_3_B phenotype. Confirmation analyses were carried out by Sanger sequencing and included samples from other family members.

**Results:** Our data revealed a novel heterozygous g.10924C>A variant on the *ABO*A*-allele located in the transcription factor binding site for RUNX1 in intron 1 (+5.8 kb site). Inheritance was shown by the results of the donor’s mother, who shared the novel variant and the anti-A specific mixed-field agglutination.

**Conclusion:** We discovered a regulatory variant in the 8-bp RUNX1 motif of *ABO*, which extends current knowledge of three other variants affecting the same motif and also leading to A_3_ or B_3_ phenotypes. Overall, long-range PCR combined with nanopore sequencing proved powerful and showed great potential as emerging strategy for resolving cases with cryptic ABO phenotypes.

## Introduction

Reduced activity of the ABO glycosyltransferase protein is in the vast majority of cases caused by single nucleotide variants (SNVs) in exonic and splice site regions of the *ABO* gene[1]. However, a particular complex phenotype of reduced activity, the so-called A_3_ or B_3_ phenotype, which presents as a mixed-field agglutination with anti-A or anti-B, can also be caused by variation in *ABO* regulatory elements located in flanking or intronic regions of the gene[2]. Standard diagnostic workflows often fail to detect these variants, as sequencing is typically limited to exons. Correctly characterizing A_3_ and B_3_ phenotypes is made even more challenging since mixed-field agglutination can also be caused by either chimerism (i.e., the presence of allogenic blood cells as a consequence of twin pregnancy[3], recent transfusion, or stem cell transplantation) or hematopoietic mosaicism[4]. To unambiguously resolve such cases, additional laborious methods like digital PCR (dPCR) or sequencing to a high read-depth using short- or long-read sequencing need to be performed.

Latest third-generation long-read sequencing by Oxford Nanopore Technologies (ONT) has the potential to resolve these complex weak ABO phenotypes. In combination with long-range PCR[5], this method enables the generation of fully resolved haplotypes of the entire gene including flanking regions. Additionally, high read-depth obtained by amplicon sequencing allows identifying cases of chimerism or mosaicism by facilitating variant calling in subclonal cell populations. Here, we describe the case of a 23-year-old female blood donor presenting with an A_3_B phenotype. Sequencing the entire *ABO* gene with ONT, we found an unknown SNV affecting the RUNX1 motif in the regulatory 5.8 kb site in *ABO* intron 1. Causality for the observed phenotype was corroborated by additional analyses in the index case as well as other family members.

## Materials and Methods

The ABO phenotype of the blood donor was determined by standard serological methods for ABH including gel cards (Grifols, Spain) and tube tests (Bio-Rad, Switzerland), as well as by anti-A_1_ and anti-H specific agglutination (Bio-Rad Seraclone, Switzerland). Genotyping of main *ABO* variants was carried out by commercially available kits (Inno-train, Germany) based on PCR with sequence-specific primers (PCR-SSP). Presence of A- and B-antigen on erythrocytes was quantified on a FACSCanto II flow cytometer (BD Biosciences, Switzerland) using monoclonal IgM antibodies (anti-A and anti-B from clones BIRMA-1 and LB-2, respectively). Potential chimerism was investigated by analysing the allelic distribution of 24 genetic variants across the genome using dPCR (Stilla, France). The entire *ABO* gene including flanking regions was amplified by two overlapping long-range PCRs (∼16 and ∼13 kb, respectively) and sequenced on part of a MinION R9.4 flowcell (ONT). Both ONT sequencing and bioinformatical analyses were carried out as described recently[5]. Nanopore sequencing results were verified by Sanger sequencing of all seven *ABO* exons as well as the identified candidate region containing the RUNX1 motif in intron 1. To elucidate a somatic versus germline origin of the identified candidate variant, we extended phenotypic and genetic analyses to the donor’s mother and brother. Sanger sequencing was limited to the candidate region in intron 1.

## Results

Genotyped as *ABO*A1* and *ABO*B* by PCR-SSP, the proband’s forward phenotyping showed a double population in the agglutination with anti-A and strong reactions with anti-B and anti-AB (Table 1). Agglutination with anti-H and anti-A_1_ was weak and absent, respectively. All observations were in agreement with an A_3_B phenotype. Flow cytometry revealed that ∼80% of erythrocytes lacked the A-antigen. No sign of chimerism was identified by dPCR.

**Table 1.** Serological and genetic results of index case, mother and brother.

|  | Serology / Forward Phenotyping |  |  |  |  | Reverse Phenotyping |  |  |  | Genetics |  |  | Phenotype |
| --- | --- | --- | --- | --- | --- | --- | --- | --- | --- | --- | --- | --- | --- |
|  | Anti-A | Anti-B | Anti-AB | Anti-H | Anti-A<sub>1</sub> | A1 | A2 | B | O | PCR-SSP | Sequencing (ONT) | Sequencing (Sanger) |  |
| **Index Case** | mf<sup>a</sup> | 4+<sup>b</sup> | 4+ | 1+ | 0 | 0 | 0 | 0 | 0 | *A1/B<sup>c</sup>* | *A1+g.10924C>A\|B.01<sup>d</sup>* | *g.10924C/A* | A<sub>3</sub>B |
| **Mother** | mf | 0 | mf | 2+ | mf | 0 | 0 | 4+ | 0 | *A1/O.01* | n/a | *g.10924C/A* | A<sub>3</sub> |
| **Brother** | 0 | 0 | 0 | n/a | n/a | 4+ | 4+ | 4+ | 0 | *O.01/O.01* | n/a | *g.10924C/C* | O |

<sup>a</sup> mixed-field
<sup>b</sup> 1-4+ refers to agglutination strength
<sup>c</sup> *ABO* allelic category is abbreviated, e.g. *A1* is short for *ABO\*A1*
<sup>d</sup> g.10924 coordinate refers to *ABO* reference sequence NG\_006669.2

To resolve the genetic basis of the observed A_3_B phenotype, we sequenced the entire *ABO* gene. Read depth of nanopore sequencing exceeded 10,000x for both PCR amplicons allowing variant calling in subclonal cell populations. Genetic variation in the overlapping region of PCR amplicons enabled read-based phasing of all genetic variants across the entire gene. As expected, one of the two *ABO* haplotypes corresponded to an *ABO*B* allele, comprising all seven *ABO*B*.*01* defining exonic SNVs. They were all heterozygous. The second haplotype was identified as an *ABO*A1* allele. No exonic variant was found that would explain the mixed-field agglutination. However, we discovered on this haplotype a novel C>A variant at NG_006669.2:g.10924 (alternatively: NM_020469.3:c.28+5871C>A), located in a known transcription factor binding site for RUNX1 in the +5.8 kb site of intron 1 (Figure 1)[6].

**Figure 1.**
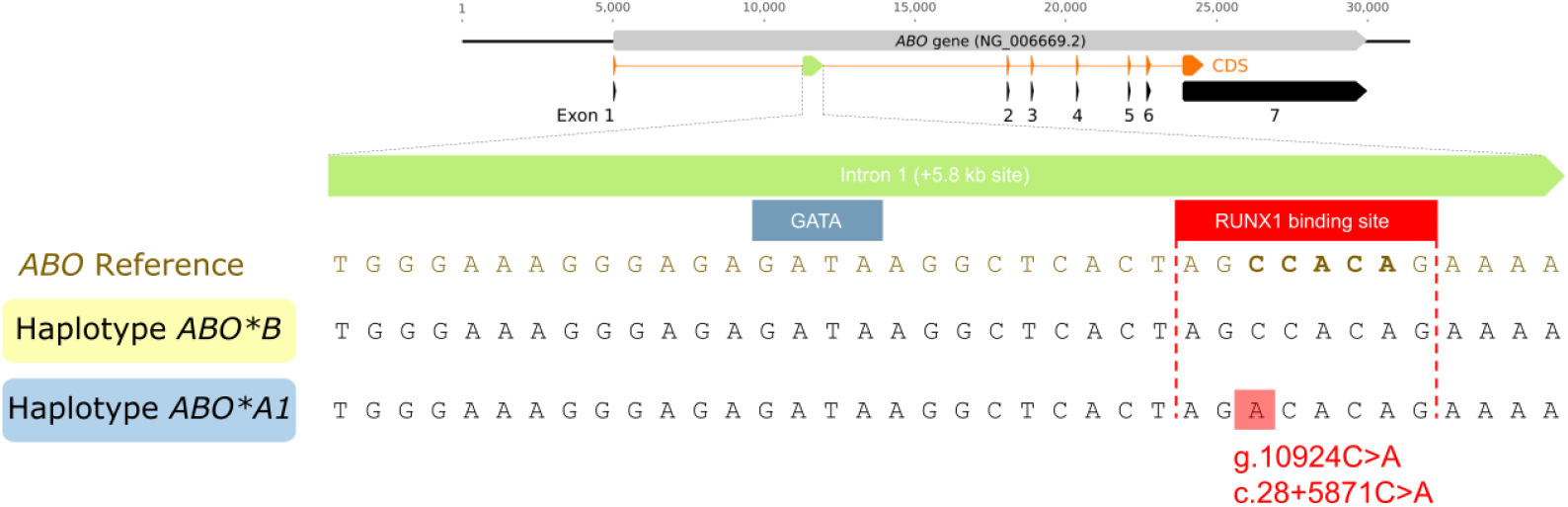
Alignment of *ABO* reference sequence (NG_006669.2) and nanopore-based maternal (blue) and paternal (yellow) haplotype sequences of index case over the regulatory 5.8 kb site in *ABO* intron. **1**. Overall genetic structure of *ABO* is provided above the alignment. The coding DNA sequence (CDS) is shown in orange. The novel regulatory variant located in the RUNX1 binding site (NG_006669.2:g.10924C>A; NM_020469.3:c.28+5871C>A) on the *ABO*A1* haplotype is highlighted with a red box. Nucleotides at the core motif for RUNX1 binding[6] (CCACA) are highlighted in bold on the reference sequence.

To further exclude the non-inherited phenomena of chimerism and mosaicism as a possible explanation for the A_3_B phenotype, we also analysed the proband’s mother and brother. The mother was genotyped *ABO*A1*/*ABO*O*.*01* by PCR-SSP and also showed a mixed-field agglutination with anti-A (Table 1). Compared to her daughter, the proportion of erythrocytes presenting A-antigen was higher (∼35%). This may support the allelic enhancement hypothesis, i.e. the assumption that the *in-trans* allele impacts the expression level of the weak allele[7, 8]. Sanger sequencing confirmed the heterozygous presence of the novel variant NG_006669.2:g.10924C>A in the RUNX1 binding site. As expected, the brother, who was genotyped as *ABO*O*.*01*/*ABO*O*.*01*, did not show any variation in the RUNX1 motif (Table 1).

## Discussion

We resolved the genetic basis of an elusive A_3_ phenotype using latest third-generating long-read sequencing. In light of the diverse sources for phenotypes of mixed-field agglutination, including genetic variants in exonic regions, regulatory elements and subclonal cell populations, we employed nanopore sequencing as our method of choice. In combination with long-range PCR, this sequencing technology allowed to comprehensively investigate the complete *ABO* gene at high read-depth as well as constructing entire *ABO* haplotypes. Our approach revealed a novel regulatory variant in the RUNX1 binding site of the *ABO* intron 1 underlying the observed A_3_ phenotype.

Typically, assessing causality for an expression-linked variant would require additional experiments like reporter gene assays or electrophoretic mobility shift assays[9]. However, in this particular case, the available evidence strongly supports the causal association. First, the presence of genetically different cell populations was excluded by dPCR and by the observed heritability of the cryptic phenotype. As a consequence, additional analyses like identifying variants in subclonal cell populations were no longer necessary. Second, beside the *ABO*B*.*01* defining SNVs, no variants were detected in the coding or splice site regions of the *ABO* gene. Third, the identified intronic C>A variant is exceedingly rare based on its absence in large databases like dbSNP, lied on the haplotype representing an *ABO*A1* allele and was inherited along with the A_3_ phenotype from the mother. Finally, and most convincingly, three SNVs targeting the core consensus sequence[6] (5’-CCACA-3’, Figure 1) of the 8-bp RUNX1 binding site had already been reported previously and had experimentally been shown to be causative for A_3_ and B_3_ phenotypes (Table 2a)[8, 10, 11]. While our understanding of the mutational consequences at positions outside the core element remains limited (Table 2a), it is noteworthy that variants in the direct vicinity have been reported to cause the same phenotype (Table 2b)[8, 12]. Interestingly, the deletion of the entire RUNX1 motif results in stronger A_m_ or B_m_ phenotypes (Table 2a)[9, 13, 14], characterised by complete absence of the respective A or B-antigen on erythrocytes. Other cell types, however, are not affected as this regulatory motif acts in an erythroid-specific manner. Accordingly, there is no formation of isoagglutinins in these phenotypes.

**Table 2.** Overview of reported functional consequences of genetic variation within and in close proximity to the *ABO* RUNX1-binding site. The RUNX1 transcription factor binding site covers positions 10922 to 10929 of the *ABO* reference sequence NG_006669.2. For a complete overview, all positions within the motif are listed, regardless of whether genetic variation has been described. All SNVs listed seem to preferentially cause phenotypes of mixed-field agglutination (A_3_, B_3_). Deletion of the entire motif results in stronger A_m_ or B_m_ phenotypes, in which none of the respective antigens is detected on erythrocytes.

| RUNX1 motif position<sup>a</sup> | Gene position<sup>b</sup> | Variation | Phenotypic consequence | N<sup>c</sup> (ethnicity) | Reference |
| --- | --- | --- | --- | --- | --- |
| **(a) Within RUNX1-binding site** |  |  |  |  |  |
| Pos 1 | g.10922A | - | - | - | - |
| Pos 2 | g.10923G | - | - | - | - |
| Pos 3* | g.10924C | C>A | A<sub>3</sub> | 2 (European) | current study |
| Pos 4* | g.10925C | C>T | B<sub>3</sub> or B<sub>w</sub> | 3 (Chinese)<br>7 (Koreans) | Ying et al.[10]<br>Yu et al.[11] |
| Pos 5* | g.10926A | A>C | B<sub>3</sub> | 1 (n/a) | Hult et al.[8] |
| Pos 6* | g.10927C | - | - | - | - |
| Pos 7* | g.10928A | A>G | B<sub>3</sub> | 4 (n/a) | Hult et al.[8] |
| Pos 8 | g.10929G | C>T | n/a | 1 (n/a) | dbSNP: rs1834858399 |
| Pos 1-8 | entire motif absent | 23 bp-del<br>3 kb-del<br>5.8 kb-del | A<sub>m</sub><br>B<sub>m</sub><br>B<sub>m</sub> | 2 (Japanese)<br>1 (Japanese)<br>111 (Japanese) | Takahashi et al.[14]<br>Sano et al.[13]<br>Sano et al.[9] |
| **(b) In proximity to RUNX1-binding site** |  |  |  |  |  |
|  | g.10930A | A>G<br>delAAAA | A<sub>3</sub><br>A<sub>3</sub> | 3 (Japanese)<br>3 (Thai) | Takahashi et al.[12]<br>Hult et al.[8] |

<sup>a</sup> all positions of the *ABO* RUNX1 binding site are listed. Nucleotides at the core consensus sequence[6] (CCACA) are highlighted by asterisks (\*)
<sup>b</sup> gene position refers to NG\_006669.2
<sup>c</sup> number of reported cases

Overall, long-read sequencing greatly simplified the discovery of the causal variant of the observed A_3_ phenotype as it enabled the efficient investigation of the complete *ABO* gene, including regulatory sites in introns and flanking regions. Such regulatory regions may often be either unknown or overlooked, and consequently not covered by standard approaches like Sanger sequencing. Of note, variants affecting the RUNX1 motif in the *ABO* gene had so far almost exclusively been reported from East Asian populations (Table 2) and were never transferred to public databases for blood group alleles[1] or for clinically relevant variant data (e.g. ClinVar). Lack of submission to appropriate databases unfortunately hampers knowledge dissemination and may foster incomprehensive investigation of such weak or cryptic phenotypes.

In summary, we detected a novel regulatory variant in the *ABO* RUNX1 binding site in intron 1 by ONT sequencing. The resulting A_3_ phenotype adds to the growing knowledge on mutational consequences in this regulatory element. More broadly, long-range PCR combined with long-read sequencing represents a promising strategy to generally resolve cases with cryptic ABO phenotypes.

## Acknowledgements

S.M. and M.P.M.-G. designed the study, supervised the data analysis and reviewed and edited the manuscript. G.A.T. and M.G. performed the experiments, analysed the data and wrote the manuscript. S.S., E.M, E.G., L.S., Y.M., N.T. and K.N. contributed to experiments. C.E. and B.M.F. contributed to the design of the study and facilitated sample acquirement. All authors read and approved the final manuscript.

We thank the proband’s mother and brother for their willingness to donate blood for research purpose.

## Data Availability Statement

A 402 bp region of the +5.8 kb site of *ABO* intron 1, causative for ABO blood group phenotype A_3_, has been submitted to GenBank (accession number OP479978). The identified variant has been submitted to ClinVar (accession number VCV001707607.1).

